# Phosphotungstic acid (PTA) preferentially binds to collagen-rich regions of porcine carotid arteries and human atherosclerotic plaques using 3D micro-computed tomography (CE-μCT)

**DOI:** 10.1101/2022.07.14.499520

**Authors:** A. Hanly, R. D Johnston, C. Lemass, A. Jose, B. Tornifoglio, C. Lally

**Author notes:** Both authors contributed equally to this work (A. Hanly, R. D Johnston).

## Abstract

**Background and aims:** Atherosclerotic plaque rupture in the carotid artery can cause small emboli to travel to cerebral arteries, causing blockages and preventing blood flow leading to stroke. Contrast enhanced micro computed tomography (CEμCT) using a novel stain, phosphotungstic acid (PTA) can provide insights into the microstructure of the vessel wall and atherosclerotic plaque, and hence their likelihood to rupture. Furthermore, it has been suggested that collagen content and orientation can be related to mechanical integrity. This study aims to build on existing literature and establish a robust and reproducible staining and imaging technique to non-destructively quantify the collagen content within arteries and plaques as an alternative to routine histology.

**Methods:** Porcine carotid arteries and human atherosclerotic plaques were stained with a concentration of 1% PTA staining solution and imaged using MicroCT to establish the in-situ architecture of the tissue and measure collagen content. A histological assessment of the collagen content was also performed from picrosirius red (PSR) staining.

**Results:** PTA stained arterial samples highlight the reproducibility of the PTA staining and MicroCT imaging technique used with a quantitative analysis showing a positive correlation between the collagen content measured from CEμCT and histology. Furthermore, collagen-rich areas can be clearly visualised in both the vessel wall and atherosclerotic plaque. 3D reconstruction was also performed showing that different layers of the vessel wall and various atherosclerotic plaque components can be differentiated using Hounsfield Unit (HU) values.

**Conclusions:** The work presented here is unique as it offers a quantitative method of segmenting the vessel wall into its individual components and non-destructively quantifying the collagen content withing these tissues, whilst also delivering a visual representation of the fibrous structure using a single contrast agent.

**Graphical Abstract:** 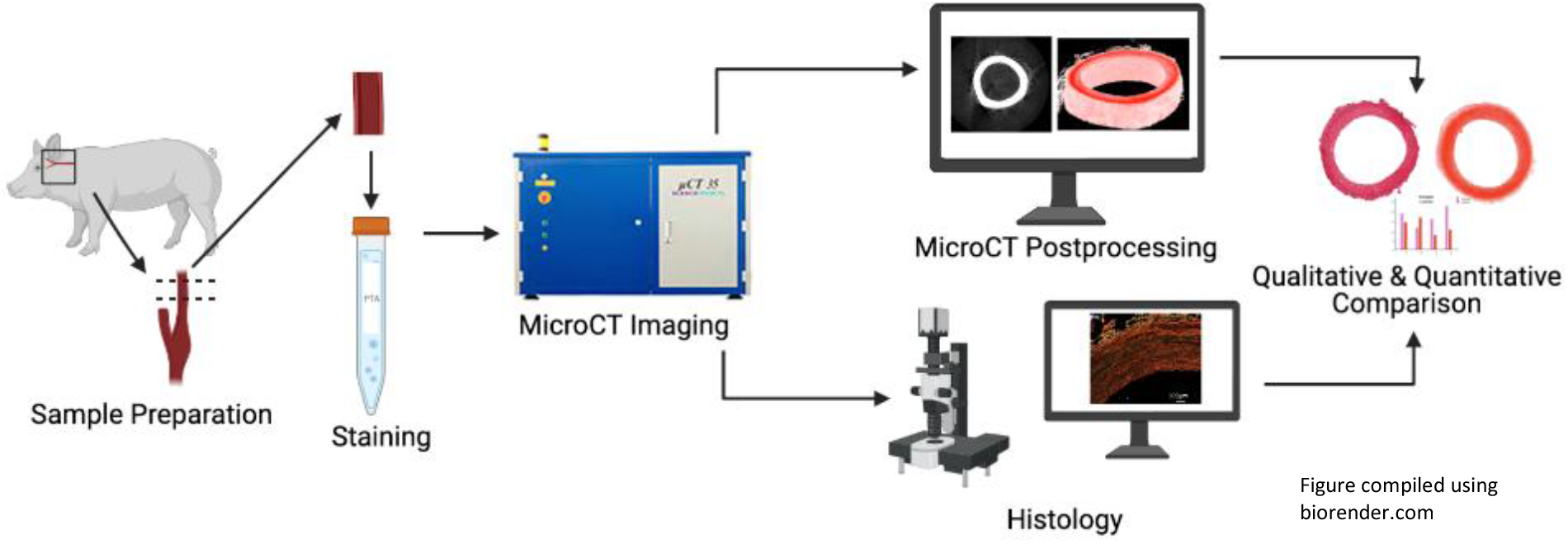

## Introduction

Plaques in the carotid artery can rupture and yield emboli that can migrate, causing blockages in smaller arteries, leading to ischemic stroke [1]. Stroke is a leading cause of death and disability in Europe. It is estimated 10% of individuals will die within 30 days of stroke onset and that over 50% of survivors will lose their independence within the first six months of recovery [2–4]. Prevention remains the best way to reduce mortality and disability as a result of stroke [5]. Collagen is known to be the major load bearing component of healthy arterial walls and atherosclerotic plaque tissues, and it is the structure and orientation of collagen within these tissues that dictates their mechanical strength [6–8]. Therefore, understanding the underlying microstructure of the arterial wall and atherosclerotic plaque tissue, along with the mechanisms of rupture, could help to identify vulnerable plaques and prevent plaque rupture. Thus, it is crucial to understand the orientation, density, and coherence of collagen fibres in both healthy and diseased states [9]. Current *in-vivo* imaging techniques remain limited in providing a detailed characterization of the vessel and atherosclerotic plaque microstructure due to resolution limitations [10] and therefore destructive *ex-vivo* techniques are used. Histology remains the gold standard for *in vitro* imaging of collagen in vascular tissue, despite its destructive nature and that it only offers a thin, two-dimensional (2D) snapshot of the tissue [11]. Optimised collagen imaging techniques that are inexpensive, fast, and reliable are of great interest, and recent works have attempted to establish new and novel imaging techniques which give a better representation of the 3D vessel microstructure [12–17].

Contrast enhanced micro computed tomography (CEμCT, MicroCT) is one such technology that offers the potential to non-destructively image the vessel wall and atherosclerotic plaque at sufficient resolution [10,13,17,18]. MicroCT is an imaging technique which can create highly specific 3D renders, depicting the internal detail of biological samples based on their interaction with X-rays [19]. Traditionally, MicroCT imaging was reserved for hard, dense tissue such as bone, but over the past decade developments in heavy element staining protocols have established it as a method to image soft tissue [20–23]. MicroCT imaging offers a unique opportunity to image the tissue microstructure at resolutions similar to histology, with the additional advantage of 3D reconstruction, and without destroying the sample of interest. Furthermore, some papers have reported reversible staining protocols, which can remove the staining solution from the tissue, leaving it entirely unaltered by the CEμCT protocol [18,24].

Recently, iodine-based stains have been used to image arteries and atherosclerotic plaques with MicroCT, and the resulting images enabled the layers of the arterial wall to be differentiated and the segmentation of calcium deposits [17,18,25]. However, none of these contrasts provide quantitative insight into the fibrous microstructure of the vessel and atherosclerotic plaque. In 2015, Nierenberger et al. [13] used a phosphotungstic acid (PTA) staining protocol, initially proposed by Metscher et al [21], to successfully image the microstructure of porcine iliac veins. Their work identified the intima, media, adventitia and connective tissue of the vein, and produced 3D renders which qualitatively showed the orientation of the fibrous structures. Based on studies that report that PTA binds preferentially to collagen [19,26,27], Nierenberger et al. concluded that it was the collagen microstructure within these vessels which was rendered in 3D. However, the work of Nierenberger et al. did not have histological controls to verify their findings, therefore further exploration of the binding element of PTA to collagen is required. Furthermore, the work of Nierenberger et al. gave no indication of collagen content within their samples and was limited to qualitatively noting the orientation of the microstructure.

Within this study, we tested the hypothesis that the staining protocols presented by Nierenberger et al. [13] could be adapted and applied to porcine arterial tissue, and to human atherosclerotic plaques. We explored whether, when coupled with MicroCT imaging, these protocols could be used to image the collagen in these tissues in 3D, and yield both qualitative and quantitative information on the collagen content within the tissues. We compared the images obtained from PTA staining and MicroCT imaging to Picrosirius red (PSR) stained histological sections from the same sample to establish if the MicroCT protocols were compatible to routine histology, and to verify if in fact the images were indicative of the underlying collagen microstructure.

## Materials and Methods

### Sample Preparation

Porcine carotid arteries of 6-month-old white pigs were obtained from an abattoir. Within 3 hours the arteries were excised from the connective tissue and frozen at a controlled rate of −1°C/min to −80°C in tissue freezing media made up of 500 mL Gibco RPMI 1640 Medium (21875034, BioSciences), 19.6 g sucrose (S0389, Sigma) and 73.3 mL of the cryoprotectant dimethylsulfoxide (PIER20688, VWR International). The samples were stored at −80°C until ready to image. For imaging, three arteries were thawed at room temperature, rinsed with PBS and any remaining connective tissue was removed using a fresh scalpel. Two 15 mm samples from each of the three vessels were cut (n=6). Each sample was fixed in 5 ml of 10% formalin for 25 hours. After fixation they were stored in 70% ethanol until staining.

### Staining

A 1% PTA solution was made up by combining 5 ml of 10% Aqueous PTA (Sigma-Aldrich, HT152) with 45 ml of 70% ethanol. Each of the six samples was submerged in 14 ml of this solution for 16 hours. For the final 30 minutes of staining, the samples were placed on a rotator at 20 RPM. Following staining all samples were dipped in 15 ml of 70% ethanol to remove excess PTA, and then stored in 5 ml of 70% ethanol until scanning.

### MicroCT

Each sample was placed in an 11.5 mm diameter holder and the holder was filled with 70% ethanol such that the sample was submerged. The sample was secured in place using a small piece of low-density polyethylene and the top of the holder was sealed with parafilm. The holder was placed in the MicroCT chamber for imaging. Samples were imaged using a SCANCO Medical AG MicroCT42 with an X-ray energy of 70 kVp and an X-ray intensity of 114 μA. The highest resolution voxel size achievable was 6 μm. The MicroCT scan parameters were based on those of Nierenberger et al. [13] and are detailed in Table 1 of the supplementary data. After scanning, samples (n=6, from 3 different vessels) were stored in 70% ethanol and transferred to histological processing.

### De-staining

PTA was removed from the samples (n=6, paired to samples still stained for each vessel) using 0.1 M NaOH, as proposed by Schmidbaur et al [24]. 0.1 M NaOH was made up by combining 0.2 g of NaOH with 50 ml of DI water. Samples (n=6) were placed in 14 ml of the NaOH solution for 16 hours. For the final 30 minutes of de-staining samples were placed on a rotator set to 20 RPM. Samples were dipped in 15 ml of 70% ethanol to remove excess NaOH and stored in 5 ml of 70% ethanol after de-staining, some were reimaged to prove the efficacy of the washing protocols (n=3) and all 6 samples were then sent to histological processing.

### Histology

Both PTA stained (n=6) and de-stained samples (n=6) were step-wise dehydrated in ethanol to xylene, embedded in paraffin wax and sliced at a 6 μm thickness using a microtome (RM-2125RT, Leica). Slices were mounted on slides and stained with Picrosirius Red (PSR). The slides were imaged using an Olympus BX41 microscope with Ocular V2.0 software. Samples were imaged at 2x and 10x magnification. Slides were imaged using both Brightfield and Polarised Light Microscopy (PLM). Two PLM images were obtained at 90° to each other and combined to visualise the birefringent collagen and determine the collagen content within the slice.

### Data Analysis

The MicroCT data was exported in DICOM image format and the images were analysed using Horos (a free and open-source code software program available from Horosproject.org and sponsored by Nimble Co LLC d/b/a Purview in Annapolis, MD USA) and OsiriX Lite (an open-source software available from osirix-viewer.com), and MATLAB (MathWorks, Cambridge, UK).

#### Quantitative Analysis

The DICOM images were read into MATLAB, and the pixel values within the image were converted to Hounsfield Units using the DICOM header information. Three regions of interest in the internal media and three regions of interest in the external media were manually drawn (as shown in Figure 4B) and the average Hounsfield Unit value in the area was obtained. This was repeated for three slices across each of the samples and averaged.

#### Vessel Segmentation

The vessel was segmented into the intima, media and adventitia based on the Hounsfield Units within each of the layers. The thresholds used for each of the layers is included in Supplementary Table 2.

#### Histological Analysis

Collagen content was calculated using the established protocol used in Johnston et al [8] and is defined as the area of collagen in the combined PLM image divided by the total tissue area in the brightfield image. Specifically, both the 10x PLM image and the corresponding Brightfield image were binarized. The collagen content in the vessel wall was calculated as the number of white pixels in a user specified ROI in the PLM image, divided by the number of white pixels in the same ROI in the brightfield image.

### Application to Human Atherosclerotic Plaques

To establish that this protocol could be adapted for thicker and more heterogeneous plaque tissues, a human plaque obtained from endarterectomy was stained in 15 ml of 1% PTA. The plaque samples were obtained from symptomatic endarterectomy patients at St. James’ hospital, Dublin. Ethical approval for obtaining these plaque specimens in this study was obtained from St. James Hospital ethical committee in compliance with the declaration of Helsinki. Atherosclerotic plaques were washed in phosphate buffered saline and then frozen as described for the porcine carotid tissue above. The plaque was fixed in 15 ml of 10% formalin. After fixation the sample was stored in 70% ethanol until staining. The sample was stained for 68 hours in total and the final 30 minutes of staining was performed on the rotator. The atherosclerotic plaque was scanned using the same scan parameters as the healthy porcine tissue, but due to the large size of the plaque the field of view was increased, such that the highest resolution possible was 8μm. The images were exported in DICOM format and analysed using Horos.

#### Statistical Analysis

Statistical analysis was performed with Prism 8 statistical software (GraphPad Software Inc., San Diego, California) via a Pearson correlation analysis. This allowed for the relationship between the Hounsfield unit measured from MicroCT to the collagen content observed in histology to be established.

## Results

### PTA Staining

While an unstained artery (termed native) showed no discernible contrast with respect to the ethanol in the MicroCT images (see Figure 1A), PTA-stained arteries could be clearly distinguished from the ethanol background (see Figure 1B, 1C 1E, 1G)

**Figure 1:**
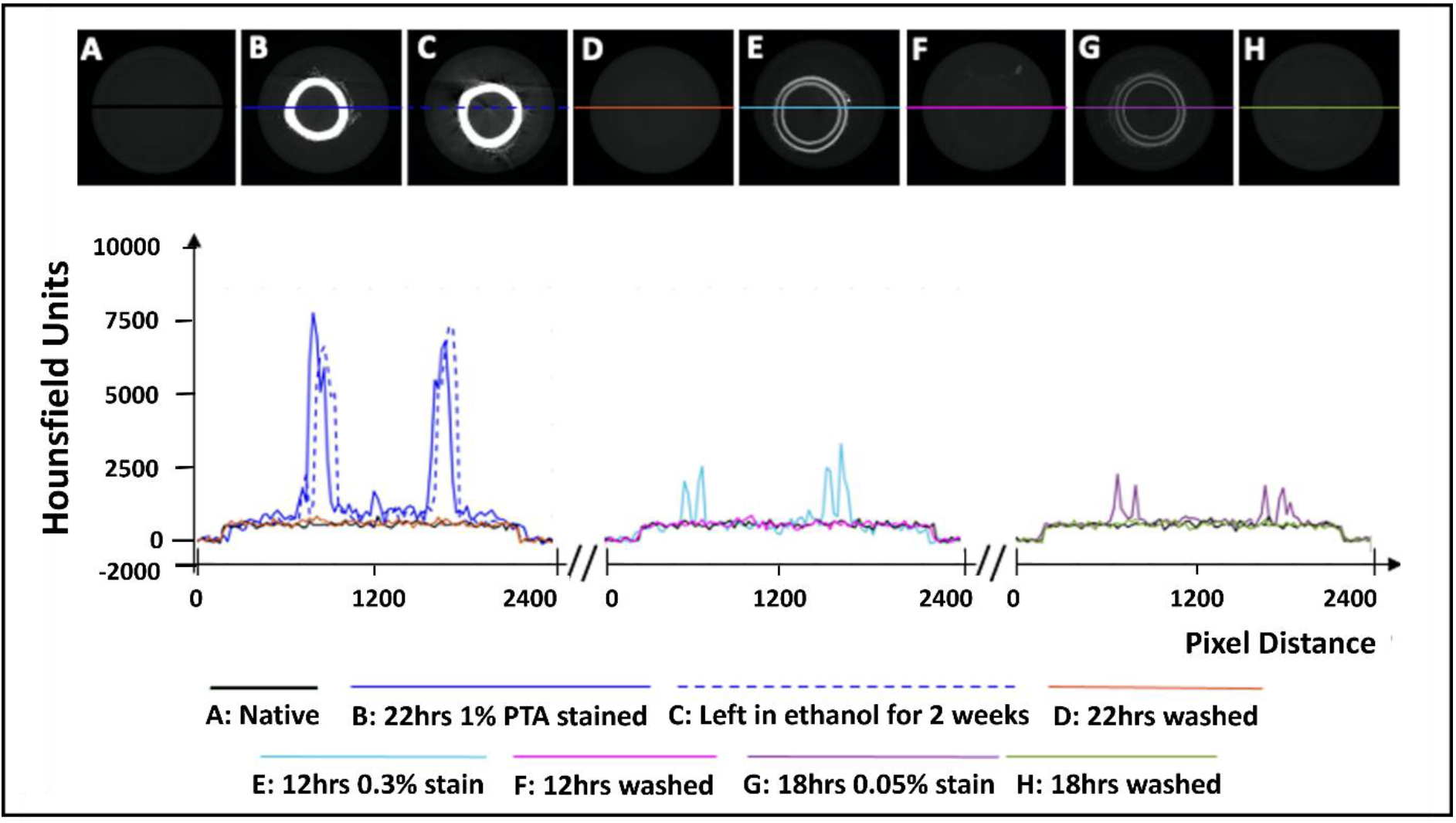
Establishment of the optimum concentration for PTA penetration through arterial tissue samples. (A) Native control (B) Stained sample for 22 hours at 1% PTA (C) Preservation in ethanol for 2 weeks (D) Removal of stain using NaOH for 22 hours (E) Stained sample for 12 hours at 0.3% PTA (F) Removal of stain using NaOH for 12 hours. (F) Stained sample for 18hrs in 0.05% stain. (G) Removal of stain using NaOH for 18 hours.

Initial exploratory work revealed that staining healthy porcine carotid with 0.3% PTA did not allow for complete penetration of the stain through the arterial wall (see Figure 1E), even after 24 hours (see Figure S1 a-c). In contrast, staining a vessel with 1% PTA allowed for full penetration of the stain through the vessel after 22 hours (see Figure 1B). Further investigation established the optimal timing for staining was 15 hours with 1% PTA concentration and this was used for the quantitative analysis (see Figures 2, 3, 4).

**Figure 2:**
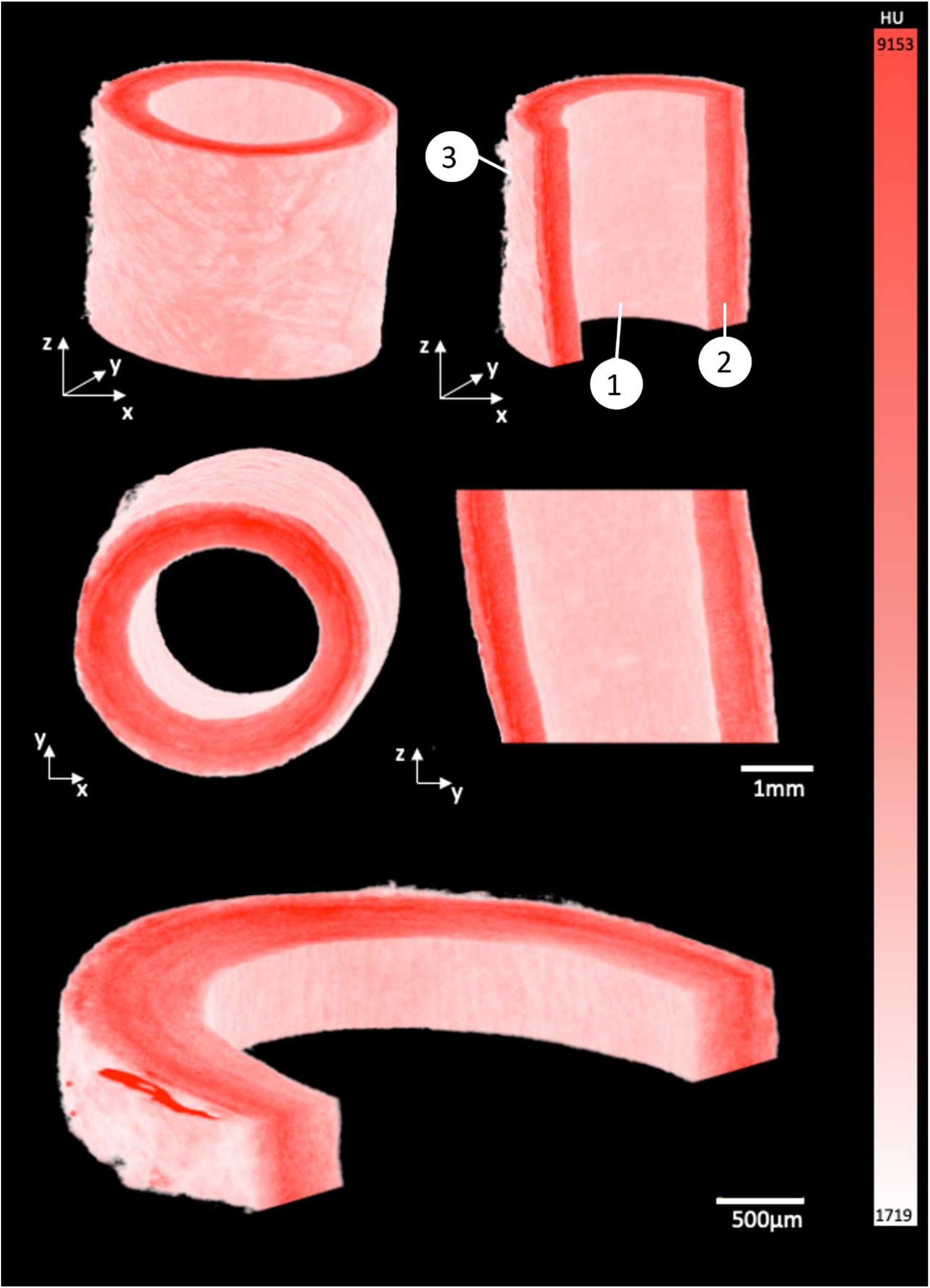
3D render of porcine carotid artery stained for 15 hours using 1% PTA. Each image shows different cross-sectional views highlighting the capability to visualize the multiple layers of the vessel wall: (1) Intimal layer, (2) Medial layer, and (3) Adventitial layer.

**Figure 3:**
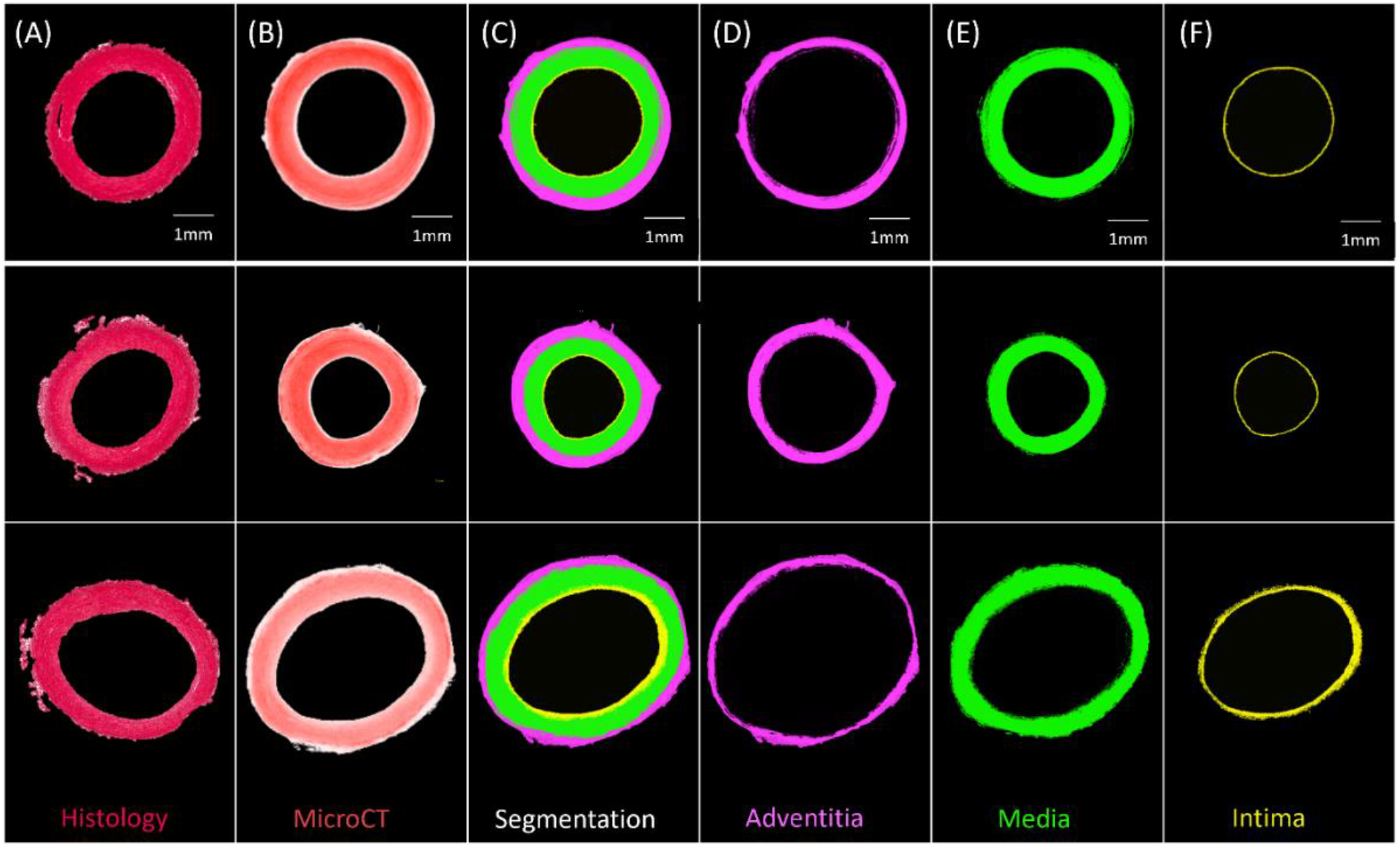
Segmentation of three PTA-stained porcine carotid artery samples into their respective layers (A) Histological PSR image, (B) PTA stained MicroCT image, (C) Segmentation, (D) Adventitia, (E) Media, and (F) Intima.

**Figure 4:**
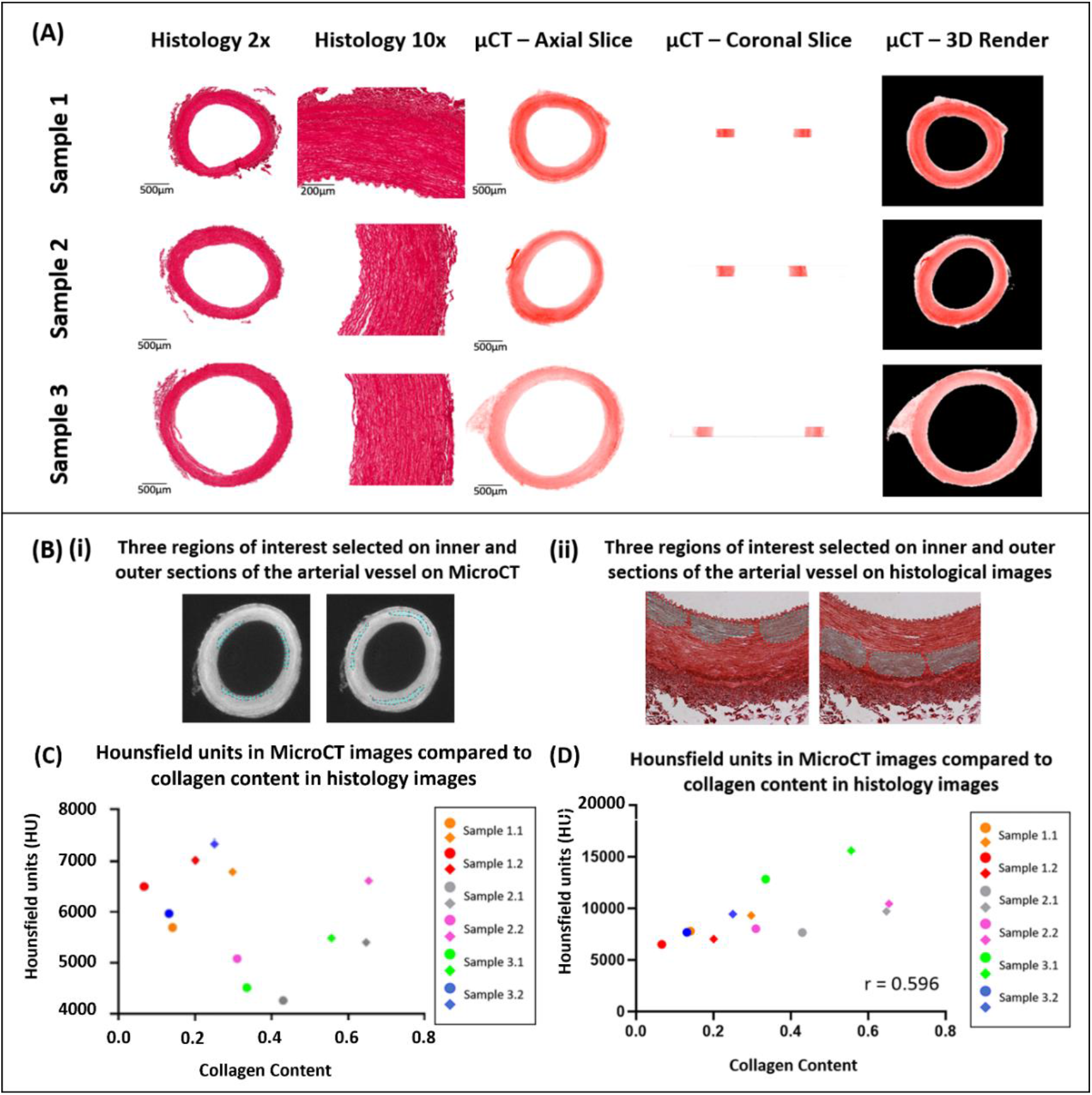
Qualitative and quantitative comparison of MicroCT to histology (A) Qualitative comparison observing the overall structure of porcine carotid arteries. (B) Quantitative comparison of the collagen content observed in porcine carotid arteries. (C) Increase in collagen content observed in different sections of vessel wall whereby the external media present higher collagen content and HU then the internal media, internal (circles) and external (diamonds) regions of interest in the media. (D) Pearson correlation between HU from MicroCT and collagen content from PS-Red images when normalised with respect to volume.

Line profiles were generated of representative cross-sections of the native, stained and de-stained arteries (see Figure 1), in order to demonstrate quantitatively the changes in radiopacity. Stronger concentrations of PTA lead to higher radiopacity across the vessel wall and higher pixel intensities, see Figure 1.

### De-staining

Exploratory work also revealed that after storage in 70% ethanol, samples stained with 1% PTA showed no discernible loss of contrast (see Figure 1C). However, when the samples were washed in 0.1M NaOH, the stain was completely removed from the sample and the pixel profile across the image returned to that of a native vessel (see Figure 1D, 1F and 1H). This was shown on all three of the samples which were re-imaged following de-staining.

### 3D Render

By using the window level tool in Osirix, a window width of 7884, and a window level of 5211 was set to ensure the samples were easily separated from the background and could be rendered in 3D, see Figure 2. The intima, media and adventitia could be distinguished in the 3D renders, and the fibrous structure within each layer was visible. Along the intima, Voxel intensity appeared to yield contrast parallel to blood flow. In the media a circumferential fibrous alignment could be seen, and the outer layer showed a loose and disorganised fibrous structure.

### Segmentation in 3D space

Samples could be quantitatively segmented into the lumen, intima, media and adventitia, based on the differences in Hounsfield Units as outlined in Supplementary Table 2.

### Qualitative Comparison of MicroCT and Histology

In comparison to conventional histology, MicroCT allowed for planar slices through the samples in the axial, sagittal and coronal planes to be viewed (see Figure 4 for examples), and for the entire artery to be rendered in 3D (see Figure 2). The MicroCT images were not prone to tissue processing defects observed in histology such as wrinkling, tearing or compression. Furthermore, MicroCT images at this resolution showed similar visualisation to 2x histology images (see Figure 4A). Within the 2D slices from the MicroCT and the histology images, the media and adventitia could be clearly identified, see Figure 4A, and in the 3D render from the MicroCT the intima, media and adventitia were clearly visible, see Figure 2. A comparison of the images obtained from MicroCT and histology is shown for three representative samples in Figure 4A.

### Quantitative Comparison of MicroCT and Histology

For each artery, it was observed that regions with higher collagen content had higher Hounsfield Units, see Figure 4C. However, across samples there was no correlation between the Hounsfield Units and the collagen content within the vessel.

It was observed that this was likely due to varying tissue volumes across the samples and the resulting variability in the intensity of the stain across different samples. Thus, the Hounsfield units were normalised with respect to volume. To do this, the volume of each tissue sample imaged was calculated on Osirix and each tissue volume was then divided by the smallest tissue volume to find a normalizing ratio. This ratio was then multiplied by the HU. Once volume was accounted for, the collagen content observed across a number of tissue samples showed a positive moderate correlation with an r = 0.596 between the histology and MicroCT as shown in Figure 4D.

### Application to Human Atherosclerotic Plaques

The staining protocol was successfully adapted to human atherosclerotic plaques. By leaving the sample in the PTA solution for a longer period of time, the samples could be imaged using the MicroCT and rendered in 3D (see Figure 5). Qualitative assessment revealed distinctive regions within the plaque, including the lumen, lipid rich cores, calcified regions and fibrous caps, as shown in Figure 5B. However, in the atherosclerotic plaque images, the fibrous structure was not as clearly delineated as it was in the images of the healthy arteries.

**Figure 5:**
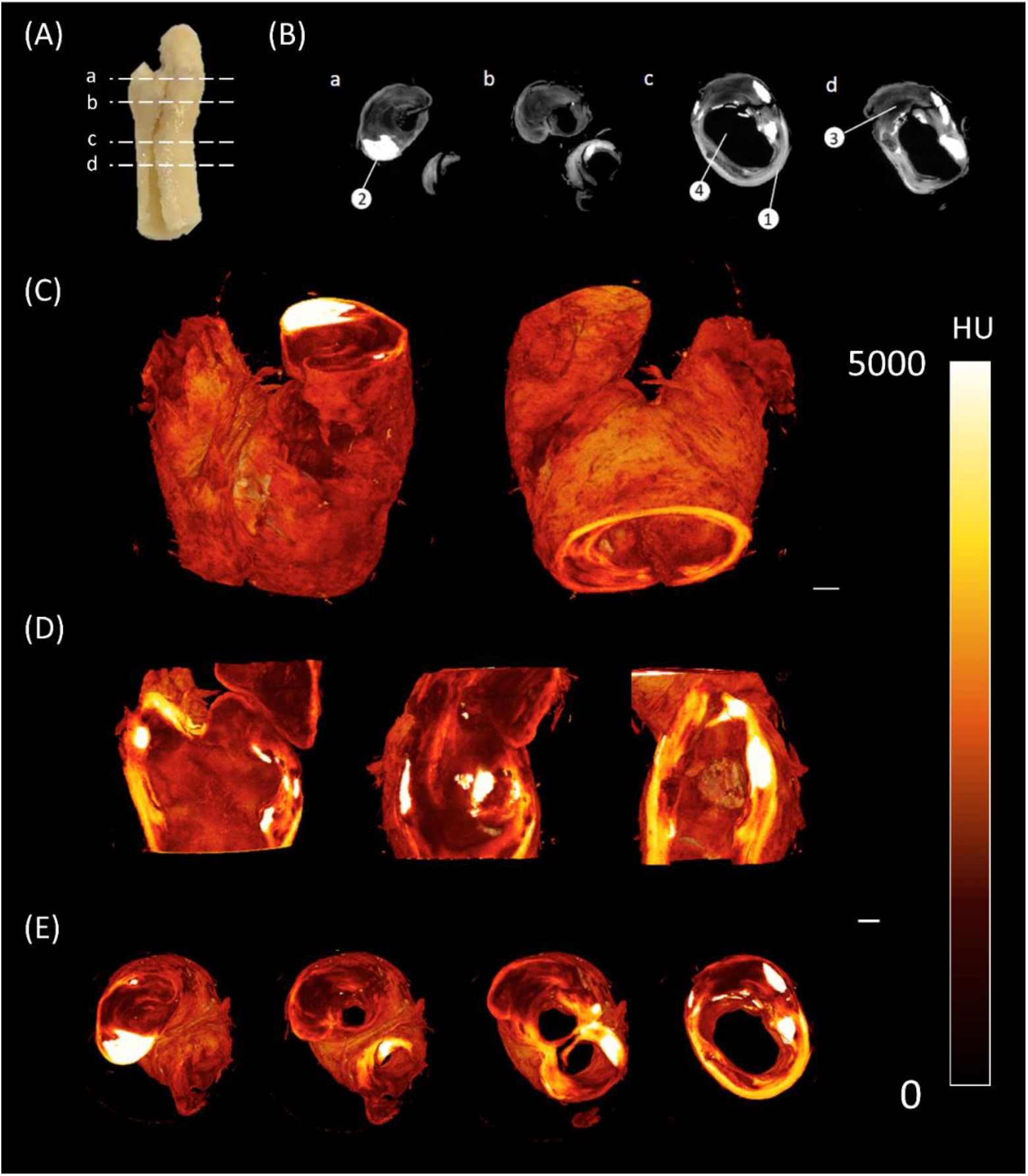
CEμCT of human atherosclerotic plaque (A) Slice locations in atherosclerotic plaque. (B) Axial cross-sections directly from MicroCT show multiple components within the tissue: (1) Outer wall of plaque, (2) Calcification, (3) Lipid, and (4) Lumen of plaque. (C) 3D render of atherosclerotic plaque. (D) Sagittal and coronal cross-sections through atherosclerotic plaque. (E) Axial cross-sections through atherosclerotic plaque. Scale bar = 1mm.

## Discussion

This data shows that MicroCT imaging, combined with PTA staining, can be used to generate high resolution 3D images of healthy and diseased arterial tissue, and has the capability to differentiate key features of the vessel wall and delineate the fibrous structure within these tissues. While CEμCT has been used to image arterial tissue in the past, the fibrous structures within the vessels were not visible. We found the protocols established by Nierenberger et al. for staining the fibrous structures in veins were not transferrable to porcine arterial tissue, as the 0.3% PTA stain repeatedly failed to penetrate through the vessel wall, even after 24 hours. There are a number of plausible reasons for this; firstly, although arteries and veins are composed of the adventitia, media and intima, the wall thickness of veins is significantly thinner than arteries, with larger lumen diameters. Whilst the variation in thickness would affect the staining time, as suggested by Pauwels et al [23], the presence of internal and external elastic laminae in arteries could also affect the diffusion capability of PTA through the tissue. Secondly, although veins and arteries are comprised of the same structural layers as mentioned previously, the size of these layers is different. In arterial tissue, the thickest layer is the tunica media which comprises a number of microstructural components such as collagen, elastin and smooth muscle cells. This layer is much thinner in the veins. The relative content of these components in the tissue are different given the different structural and functional requirements of veins and arteries. Using a higher concentration of PTA, we repeatedly saw full penetration of the stain through the vessel wall, provided the samples were left in the stain for 15 hours or more. A 1% PTA solution has previously been used by to stain cartilage samples for MicroCT imaging, and a 10% PTA solution has been used to stain mouse plaque samples [28], but to the author’s knowledge this is the first time an established PTA protocol has been used to stain porcine or human arterial tissue with quantitative analysis of the collagen content within these samples.

In the healthy porcine arteries, at an isotropic voxel size of 6μm the fibrous structure within the vessel wall was visible, and changes in the fibre orientation between the intima, media and adventitia could be visualized. The circumferential fibre alignment seen in the medial layer of the 3D render and the more disorganised fibre alignment in the adventitia agree with what is widely reported in literature to be the fibrous structure of arteries [15,29–31]. The internal layer of the vessel wall also showed elongated parallel grooves, which corresponds to the parallel alignment of this layer with the blood flow in the vessel [15,29–31]. Given the 6μm voxel size used in the current study, the images were not quite as high resolution as those in the Nierenberger et al. study on porcine iliac veins, where the voxel size was 1μm.

The vessels imaged using MicroCT for this study were also imaged by histology, which further establishes CEμCT as an imaging modality which is complementary to conventional histology. This work has also shown that MicroCT may be an appropriate alternative to histology in some cases. Images from the MicroCT at this resolution are similar to 2x histology images, see Figure 4, and MicroCT imaging presents its own unique advantages over histology. MicroCT is less tedious and faster than conventional histology, preserves tissue integrity and enables rapid 3D visualisation of the tissue. While 3D histopathology methods have been developed, they are slow and intricate, and often rely on a very high number of slides within a z-stack [14,32]. The isotropic voxel size for MicroCT imaging means the resolution in the z-direction is equal to that in the x and y planes. With MicroCT imaging, cross-sectional images could be obtained in the coronal, sagittal and axial planes at the same resolution, on a single dataset, something that is impossible with histology.

Using Hounsfield units, the layers of the arterial wall could be reproducibly and quantitatively segmented in a manner which shows good agreement with corresponding histology images. Nierenberger et al. [13] defined layers of the vein wall but based on qualitative assessment and with no histological analysis to confirm their interpretation. Robinson et al. and Self et al. managed to segment layers of the vessel wall in iodine and iohexol stained plaque tissue, and a similar segmentation has been obtained here for healthy PTA-stained vessels [17,18,25]. In fact, this work advances the work of Robinson et al. [17], because here the boundary of every layer of the arterial wall was defined quantitatively, whereas in the work of Robinson et al. the media and the intima were manually segmented. In the work of Self et al. [25] it is stated that the vessel layers were segmented based on the changes in grey levels across the vessel wall, but the authors provide no further detail which could be used for comparison to the methods here. They do however state their method is “subjective in nature”. Thus, the method of segmentation provided in this paper offers a significant improvement where provided the stain concentration and imaging parameters are kept constant, the method can be objectively and repeatably applied to multiple vessels.

Samples which were washed in 0.1M NaOH lost the contrast generated by PTA and had pixel profiles like that of unstained arteries (Figure 1). It was necessary to use a specialized washing solution because little loss of contrast was observed in samples stored in 70% ethanol for two weeks, indicating the stain remains stable in the sample over time, which has been reported previously [13,21]. The washing protocols build on the work of Schmidbaur et al [24]., who showed 0.3% PTA could be washed from preserved Syllis Armillaris using 0.1M NaOH. In their study the PTA stain was removed from two out of three 0.3 mm samples after six hours. In this study, stain removal was achieved in all three re-imaged vessels, in which the wall thickness ranged from 0.8 – 1.2 mm. By matching both the wash volume and wash time to the staining volume and time, the 1% PTA was removed.

Quantitative analysis showed a moderate positive correlation between the MicroCT Hounsfield unit and measured collagen content from histology as shown by Figure 4C, with an r value of 0.596 once volume was accounted for. However, it was also shown that within blood vessels, the PTA concentration was higher in the more collagen rich external media than the internal media, suggesting that PTA is binding preferentially to collagen and is visualised in the MicroCT. Furthermore, it is shown that higher collagen content is observed in the medial layer of the arterial wall when compared to intima and adventitia and this is also expected as its observed in the literature [33]. This indicates that CEμCT combined with PTA staining could be used to identify more collagen rich regions within samples. To establish a more quantitative metric and compare the collagen content across multiple samples, a normalising ratio with respect to volume would be required either for the staining solution or during post-processing.

Using the amended protocol for PTA staining of atherosclerotic plaque tissue, several tissue components could be clearly visualised and quantified using MicroCT. However, detail on the microstructural level is not the same as observed in the porcine arterial vessels due to a different resolution being used. The reason for this lower resolution is due to the larger FOV size that is required. As the voxel size for imaging the plaque was 8μm, it was unlikely that individual collagen fibres would be observed. Nonetheless it has been shown that the staining protocols are transferrable to human plaque tissue and improved image resolution will be further explored in future work.

A number of limitations can be noted in this study, firstly, the selectivity of the binding of PTA to collagen has long been debated in literature [26,34]. Collagen degradation has a significant effect on the binding of PTA in articular cartilage tissue suggesting its specificity of binding to collagen [27]; however, this has not been established in arterial tissue. While this research indicates PTA binds more intensely to collagen rich areas, future work should work to further establish the specificity of the stain. Furthermore, MicroCT is a non-destructive imaging technique that preserves the mechanical properties of the vessel; however, no mechanical testing has been performed in this study. A focus for future research will be to investigate the staining of fresh arterial samples that allow for mechanical tests to be performed, similar to Brunet et al [35]. Lastly, due to resolution constraints, the maximum resolution that could be achieved was 6μm. Given that collagen fibres are approximately the same size as the resolution set and that collagen fibre diameters can vary throughout the thickness of the vessel wall [36–38], the resolution is insufficient to fully delineate fibres and perform fibre tracking to establish fibre orientation similar to diffusion tensor imaging protocols [14,39].

## Conclusion

This work supports the use of MicroCT imaging combined with PTA staining as a novel and robust protocol for assessing the structure and collagen content of healthy arterial tissue and has established that the methodology is adaptable to human atherosclerotic plaque. We have built on the findings of other key studies in the field. With one singular stain, we have achieved similar results in arteries that Nierenberger et al. [13] obtained by imaging the fibrous structure of venous walls, demonstrated a substantial improvement on the work of Robinson et al. [17], who based on histology segmented multiple layers of the vessel wall using iodine and MicroCT, and confirmed the results of Self et al. [18] whereby we have shown that the stain can be washed from the arterial tissue samples after scanning. Furthermore, this study is first to use MicroCT imaging to analyse the collagen content in arterial tissue and to include a quantitative analysis of CEμCT imaging of blood vessels. Characterising the collagen content in intact vessels in 3D will improve our understanding of the role of collagen in the progression of atherosclerotic diseases, which in turn may lead to better clinical indicators of plaque stenosis.

## Acknowledgements

Research was supported by the European Research Council (ERC) under the European Union’s Horizon 2020 research innovation programme (Grant Agreement No. 637674)

