## Supplementary Document for "Phosphotungstic acid (PTA) preferentially binds to collagen-rich regions of porcine carotid arteries and human atherosclerotic plaques using 3D micro-computed tomography (CE-μCT)"

Supplementary Information

*Supplementary table 1: MicroCT scanning parameters used for acquisition of data using Scanco Medical system.*

| MicroCT Details | SCANCO MEDICAL AG (MicroCT) |
|---|---|
| **Porcine Carotid Artery** | |
| X-ray Energy (kVp) | 70 |
| X-ray Intensity (μA) | 114 |
| Calibration | 7: 70kVp, BH: 1200mg, HA/ccm, Scaling 4096 |
| Integration Time (ms) | 300 |
| Average Data | 3 |
| Voxel Size (μm) | 6 |
| **Atherosclerotic Plaque** | |
| Voxel Size (μm) | 8 |

*Supplementary table 2: Definition for each of the layers used in vessel segmentation. Each ROI is set after attributing the following thresholds in the MicroCT images.*

| Regions of interest | Lower Threshold (HU) | Higher Threshold (HU) |
|---|---|---|
| Lumen | -1024 | 999 |
| Intima | 1000 | 3000 |
| Media | 7000 | 15000 |
| Adventitia | 3000 | 7000 |
| Connective Tissue | 1000 | 3000 |

The thresholds were set as follows; the lumen, media and connective tissue were defined using the 'grow region of interest" tool and the thresholds below in Horos. The outer media was set in the same way. The inner media was then defined as the area between the intima and the outer media, and the adventitia as the area between the connective tissue and the outer media.

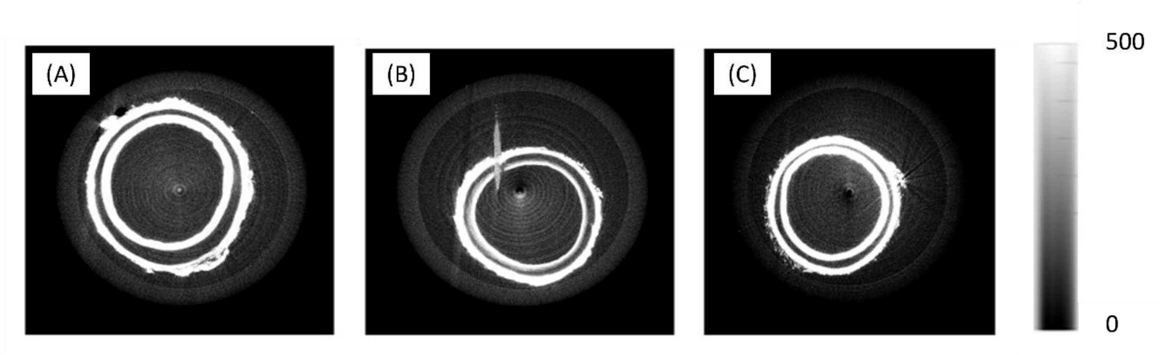

Figure S1: (A) A vessel stained for 12 hours in 0.3% PTA and finished on the rotator (B) A vessel stained for 24 hours and finished on the rotator (note the mark in the vessel is a cactus spine which was used as part of initial exploratory studies as a place marker) (C) A vessel stained for 12 hours while being rotated continuously.

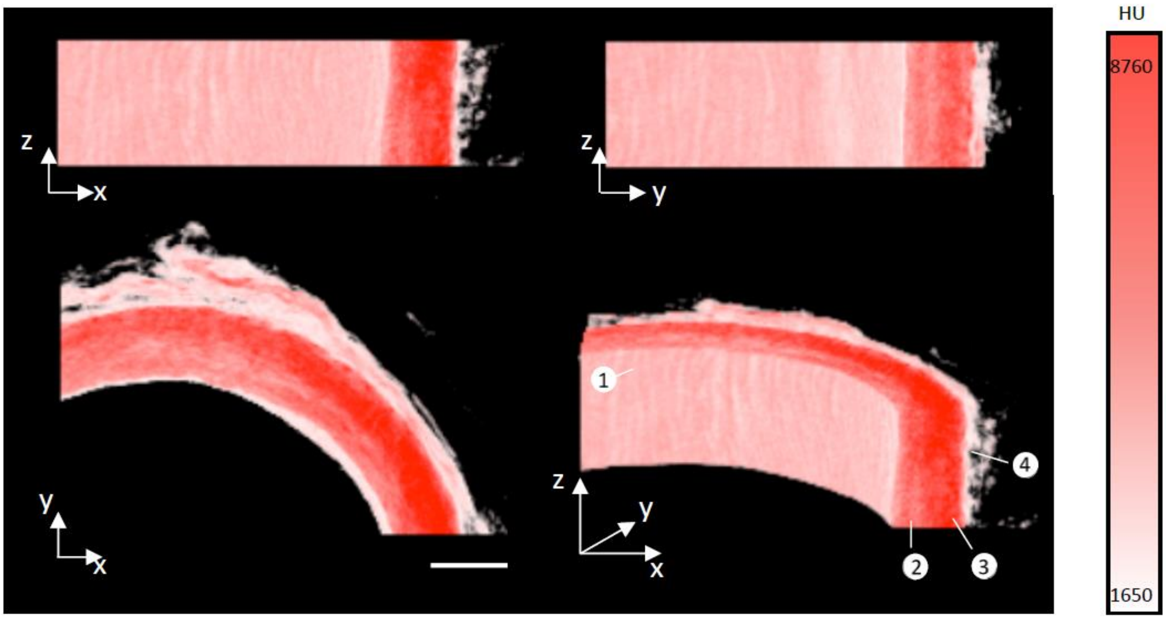

Figure S2: An enlarged segment looking at three orthogonal views and the 3D render of porcine arterial vessel imaged in this study. The (1) intima, (2) inner media, (3) outer media and (4) adventitia can be visually distinguished. Scale bar = 1 mm, HU = Hounsfield units.

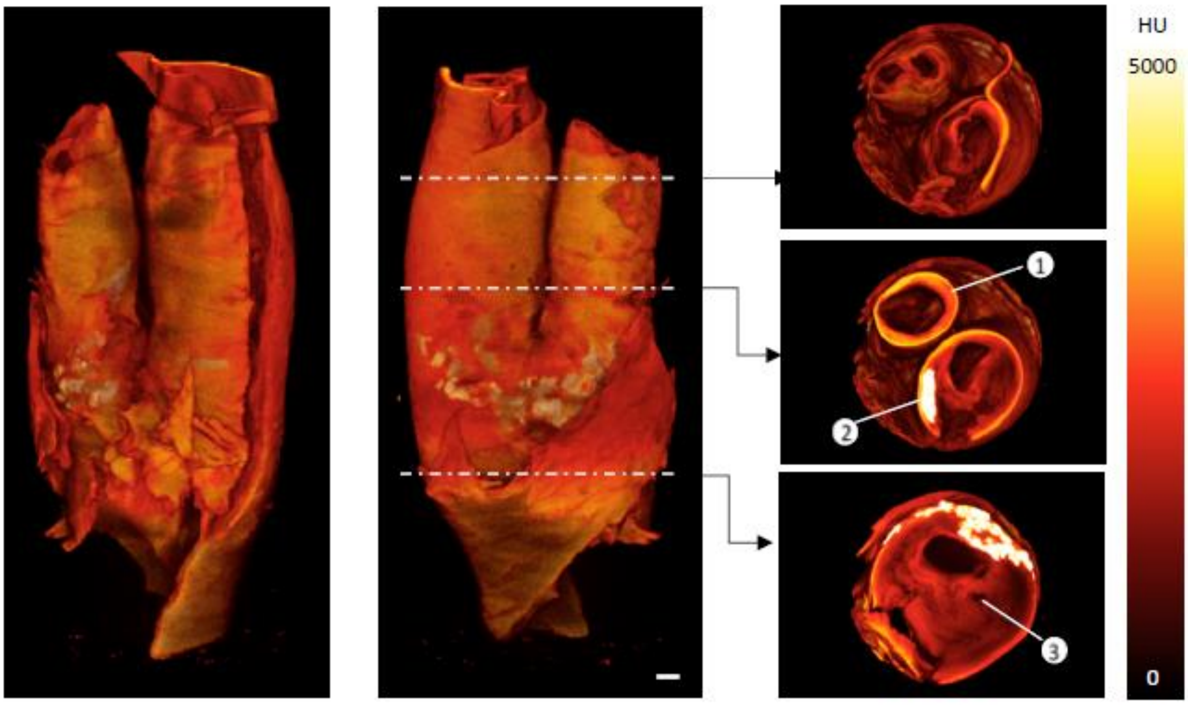

Figure S3: Second atherosclerotic plaque sample imaged in this study. The plaque geometry and internal structure can be visualized. (1) Dense fibrous tissue, (2) calcifications and (3) lipid rich cores can be identified in the 3D render. Scale bar = 1 mm, HU= Hounsfield units.
